# Proteomic analysis of the pyrenoid-traversing membranes of *Chlamydomonas reinhardtii* reveals novel components

**DOI:** 10.1101/2024.10.28.620638

**Authors:** Eric Franklin, Lianyong Wang, Edward Renne Cruz, Keenan Duggal, Sabrina L. Ergun, Aastha Garde, Alice Lunardon, Weronika Patena, Cole Pacini, Martin C. Jonikas

## Abstract

- Pyrenoids are algal CO_2_-fixing organelles that mediate approximately one-third of global carbon fixation. Most pyrenoids are traversed by membranes that are thought to supply them with concentrated CO_2_. Despite the critical nature of these membranes for pyrenoid function, they are poorly understood, with few protein components known in any species.
- Here, we identify protein components of the pyrenoid-traversing membranes from the leading model alga, *Chlamydomonas reinhardtii,* by affinity purification and mass spectrometry of membrane fragments. Our proteome includes previously-known proteins as well as novel candidates.
- We further characterize two of the novel pyrenoid-traversing membrane-resident proteins: Cre10.g452250, which we name Pyrenoid Membrane Enriched 1 (PME1), and Cre02.g143550, also known as Low-CO_2_-Induced 16 (LCI16). We confirm the pyrenoid-traversing membrane localization of LCI16 and observe that PME1 and LCI16 physically interact. We find that neither protein is required for normal membrane morphology or growth under CO_2_-limiting conditions, but that both mutants show a similar proteomic profile to those of established pyrenoid mutants.
- Taken together, our study identifies the proteome of the pyrenoid-traversing membranes and initiates the characterization of a novel pyrenoid-traversing membrane complex, building toward a mechanistic understanding of the pyrenoid.

## Introduction

Globally, photosynthetic organisms fix over 100 billion metric tons of carbon each year via the conserved enzyme Ribulose-1,5-bisphosphate carboxylase/oxygenase (Rubisco), thought to be the most abundant protein in the world (Ellis, 1979; Field *et al*., 1998; Beardall & Raven, 2020). Despite its importance to the biosphere, Rubisco runs relatively slowly for an enzyme in central carbon metabolism and is prone to catalyzing oxygenation, a wasteful reaction with oxygen (Flamholz *et al*., 2019). Some organisms overcome these limitations through the use of CO_2_-concentrating mechanisms (CCMs). These mechanisms improve the efficiency of carbon fixation by concentrating CO_2_ locally around Rubisco to increase its rate of carboxylation and decrease unwanted oxygenation (Hatch, 1987; Lüttge, 2004; Sage *et al*., 2012; Raven *et al*., 2017).

Eukaryotic algae, which mediate most of the CO_2_ assimilation in the oceans (Mackinder, 2017), nearly all use a CCM based on a chloroplast-localized organelle called the pyrenoid (Wang *et al*., 2015; Beardall & Raven, 2020; Meyer *et al*., 2020a; Barrett *et al*., 2021; He *et al*., 2023). A pyrenoid is formed by tightly clustering the cell’s Rubisco into a biomolecular condensate (Freeman Rosenzweig *et al*., 2017). Concentrated CO_2_ is released inside this condensate, increasing the carboxylation rate of Rubisco and decreasing its oxygenation. Despite their importance to the global carbon cycle, pyrenoids remain understudied and poorly understood at a molecular level.

Because pyrenoids evolved many times across independent algal lineages (Meyer *et al*., 2020a; Barrett *et al*., 2021; He *et al*., 2023), many of the specific genes involved in CCM function differ between phylogenetic groups. However, we expect that common principles of biogenesis and function underlie these convergently-evolved structures. Our approach to uncovering these principles is to study the leading model alga, *Chlamydomonas reinhardtii* (*C. reinhardtii* hereafter), whose pyrenoid is currently best understood. The existence of an extensive experimental toolbox in this organism—including well-defined genetic and proteomic protocols and a library of knockout mutants—makes it a powerful platform for the discovery and characterization of novel pyrenoid components.

The *C. reinhardtii* pyrenoid consists of a phase-separated matrix containing tightly-packed Rubisco (Freeman Rosenzweig *et al*., 2017), and is surrounded by a starch sheath that appears to limit CO_2_ leakage and is required for normal localization of CCM components (Toyokawa *et al*., 2020; Fei *et al*., 2022). Traversing the matrix are specialized membranes that are thought to mediate the delivery of concentrated CO_2_ to the matrix (Raven, 1997; Karlsson *et al*., 1998; Sinetova *et al*., 2012; Fei *et al*., 2022).

In *C. reinhardtii*, the pyrenoid-traversing membranes form a network of tubules that is contiguous with but ultrastructurally distinct from the photosynthetic thylakoid membranes (Engel *et al*., 2015). Tubules are constitutively present, including during cell division (Goodenough, 1970), in non-CCM-requiring growth conditions (Ma *et al*., 2011; Caspari *et al*., 2017), and in mutants deficient in pyrenoid matrix formation (Goodenough & Levine, 1970; Ma *et al*., 2011; Mackinder *et al*., 2016; Caspari *et al*., 2017; He *et al*., 2020). The tubules are structurally complex, being composed of a central reticulated region with narrow tubules interconnected by three-way junctions, larger cylindrical tubules which extend radially out toward the pyrenoid periphery, and smaller minitubules within the cylindrical tubules that also appear to be made of membranes (Engel *et al*., 2015). The tubule network is dynamic, with tubule quantity and diameter varying under different growth conditions (Rawat *et al*., 1996). Given the presence of the tubules in the absence of a pyrenoid matrix, it has been proposed that they play an organizational role in pyrenoid nucleation or localization (Meyer *et al*., 2020b), while their presence in non-CCM-requiring growth conditions suggests possible additional functions beyond CO_2_ delivery.

To date, several tubule-localized proteins have been identified with varying degrees of confidence. The best-established proteins that localize specifically to the tubules are Carbonic AnHydrase 3 (CAH3; Cre09.g415700), which is thought to deliver CO_2_ to the pyrenoid (Karlsson *et al*., 1998; Sinetova *et al*., 2012); MIssing THylakoids 1 (MITH1; Cre06.g259100), which is necessary for producing pyrenoid-traversing membranes (Hennacy *et al*., 2024); and Rubisco-Binding Membrane Proteins 1 and 2 (RBMP1, Cre06.g261750; RBMP2, Cre09.g416850) (Meyer *et al*., 2020b). The functions of RBMP1 and RBMP2 are not fully understood, though RBMP1, also known as BST4, shares homology with the thylakoid-localized putative bicarbonate transporters BST1-3 (Mukherjee *et al*., 2019) and impacts tubule lumen pH during dark-to-light transition, possibly by mediating bicarbonate transport across the tubule membrane (Adler *et al*., 2024).

Multiple strategies have been employed to isolate and identify *C. reinhardtii* pyrenoid proteins, including fractionation of whole pyrenoids (Mackinder *et al*., 2016; Zhan *et al*., 2018), systematic protein localization and affinity purification of protein complexes using known pyrenoid components as baits (Mackinder, 2017; Wang *et al*., 2023), and proximity labeling of pyrenoid proteins using TurboID (Lau *et al*., 2023). However, none of these methods were explicitly focused on the pyrenoid tubules.

Here, we identify tubule-localized proteins by developing a complementary approach that relies on enriching fragments of pyrenoid-traversing membranes. This approach adapts an existing affinity purification method (Wang *et al*., 2023) to isolate membrane fragments rather than protein complexes, similar to previous methods for purifying intact organelles and membrane fractions (Zhang *et al*., 2007; Chen *et al*., 2016; Yao *et al*., 2018; Chen *et al*., 2022). We show that our method successfully recovers known tubule proteins and identifies novel candidates. We further validate and characterize two of the top candidates. By expanding the known pyrenoid tubule proteome, this work contributes to paving the way toward a mechanistic understanding of pyrenoid function.

## Materials and Methods

### Strains and Culture Conditions

All insertional mutants were obtained from the CLiP collection at the Chlamydomonas Resource Center (Li *et al*., 2019) or from the recently-reported CLiP2 mutant collection (Lunardon *et al*., 2024). Mutant IDs for these strains and their wild-type background strains can be found in Table **1** below. All strains generated in this work were deposited to the Chlamydomonas Resource Center (https://chlamycollection.org).

**Table 1.** Strains used in this work.

| Chlamydomonas Resource Center ID | Strain Description | Mating Type | Source | Antibiotic Resistance |
| --- | --- | --- | --- | --- |
| CC-4533 | Wild type<br>CMJ030 | *mt(-)* | Wild type; parent strain to CLiP library (Zhang *et al.*, 2014) | none |
| CC-5415 | Wild type | *mt(+)* | Wild type; parent strain to *lci16-2* and *lci16-3* (Lunardon *et al.*, 2024) | none |
| CC-5646 | CMJ030;RBMP1-Venus-3×FLAG | *mt(-)* | Previously published | Paromomycin |
| CC-5647 | CMJ030;RBMP2-Venus-3×FLAG | *mt(-)* | Previously published | Paromomycin |
| CSI_LW07G9 | CMJ030;CPLD9-Venus-3×FLAG | *mt(-)* | CSI collection (Meyer *et al.*, 2020b; Wang *et al.*, 2023) | Paromomycin |
| CSI_LW01C3 | CMJ030;CGL129-Venus-3×FLAG | *mt(-)* | CSI collection (Wang *et al.*, 2023) | Paromomycin |
| CC-6234 | CMJ030;LCI16-Venus-3×FLAG | *mt(-)* | CMJ030 transformed with pLM005-LCI16-Venus-3×FLAG | Paromomycin |
| LMJ.RY0402.141937 | *pme1-1* | *mt(-)* | CLiP library (Li *et al.*, 2019) | Paromomycin |
| LMJ.RY0402.168140 | *pme1-2* | *mt(-)* | CLiP library (Li *et al.*, 2019) | Paromomycin |
| LMJ.RY0402.236076 | *pme1-3* | *mt(-)* | CLiP library (Li *et al.*, 2019) | Paromomycin |
| LMJ.RY0402.215307 | *lci16-1* | *mt(-)* | CLiP library (Li *et al.*, 2019) | Paromomycin |
| CLIP2.000095 | *lci16-2* | *mt(+)* | CLiP2 library (Lunardon *et al.*, 2024) | Hygromycin |
| CLIP2.000923 | *lci16-3* | *mt(+)* | CLiP2 library (Lunardon *et al.*, 2024) | Hygromycin |

Unless otherwise noted, cultures were grown in TP media to a concentration of ∼1-2×10^6^ cells mL^-1^ prior to experiments. Unless otherwise noted, cultures were grown in low CO_2_ (air, ∼0.04% v/v CO_2_) and 150 μmol ⋅ m^−2^ ⋅ s^−1^ photons.

### Membrane Affinity Purification and Mass Spectrometry

Detergent-based affinity purification and mass spectrometry of RBMP1-Venus-3×FLAG, RBMP2-Venus-3×FLAG, and LCI16-Venus-3×FLAG were performed as described previously (Wang *et al*., 2023).

For the membrane affinity purification, our objective was to isolate fragments of membranes of interest. We sought to achieve this by choosing baits localized to our membranes of interest and performing a standard affinity purification, as described in (Wang *et al*., 2023), but with the membrane disruption step changed from dissolution with detergent to shearing with sonication. We used two tubule bait proteins (RBMP1 and RBMP2) and two thylakoid bait proteins (CPLD9, Cre01.g005001; CGL129, Cre05.g233950) (Meyer *et al*., 2020b; Wang *et al*., 2023). Each bait protein was expressed in an independent strain in the CMJ030 (CC-5433) wild-type background. We performed three iterations of this membrane affinity purification experiment, each with slight differences meant to better enrich membranes or better detect the proteins enriched in the purification.

The membrane affinity purification was based on the protocol previously described in (Wang *et al*., 2023). Each strain expressing a FLAG-tagged protein was pre-cultured in 5 mL TAP media + 2 μg/mL paromomycin under ∼100-150 μmol ⋅ m^−2^ ⋅ s^−1^ photons until reaching a concentration of ∼2-4×10^6^ cells mL^-1^. 1 mL of each of these cultures was used to inoculate 50 mL of TAP + 2 μg/mL paromomycin and grown again under ∼100-150 μmol ⋅ m^−2^ ⋅ s^−1^ photons until reaching a concentration of ∼2-4×10^6^ cells mL^-1^. These cultures were then centrifuged at 1,000×g for 5 minutes, resuspended in 50 mL TP + 2 μg/mL paromomycin, and transferred to a 1L bottle containing 800 mL of TP + 2 μg/mL paromomycin. These cultures were bubbled with air at ∼150-200 μmol ⋅ m^−2^ ⋅ s^−1^ photons until reaching a density of ∼2-4×10^6^ cells mL^-1^. Cells were harvested by centrifugation at 4°C for 10 minutes at 3,000×g, washed in 40 mL of ice-cold 1×IP buffer (25 mM HEPES, 25 mM KOAc, 1 mM Mg(OAc)_2_, 0.5 mM CaCl_2_, 100 mM Sorbitol, 1mM NaF, 0.3 mM Na_3_VO_4_, and cOmplete EDTA-free protease inhibitor (1 tablet/500 mL)) then pelleted at 4°C for 10 minutes at >3,000×g. Importantly, to keep membranes intact, our IP buffer did not include digitonin, in contrast to the IP buffer used in Wang, et al. Pellets were weighed and resuspended in a 1:1 (v/w) ratio of ice-cold 2×IP buffer, and the resulting cell suspension was then dripped into liquid nitrogen to form small popcorn pellets which were stored at −80°C. Cells were broken open by grinding for 20 minutes with a Cryomiller under liquid nitrogen temperatures, and the resulting powder was defrosted on ice for at least 45 minutes and dounced 20 times in a pre-chilled Kontes Glass Col. Duall #22 Douncer (Kimble). Rather than dissolving membranes using IP buffer with digitonin and centrifuging at 12,700×g, as in the Wang et al. protocol, membranes were instead disrupted by sonicating with a probe sonicator (QSonica) in 20 cycles of (5 x 1 sec. pulse at 40% amplitude, 30 sec. rest) in a microcentrifuge tube on ice. Following sonication, lysate was clarified by centrifugation at 1,000×g for 30 minutes at 4°C to remove cellular debris. 750 μL of the supernatant containing the membrane fragments was incubated with Anti-FLAG M2 Magnetic Beads (Sigma) that had been pre-washed twice with 1 mL 1×IP buffer and thrice with 1×IP buffer plus 0.1% digitonin (twice with 1 mL and once with 150 μL). The membrane fragment–containing lysate was nutated with the Anti-FLAG beads for 1.5 hours at 4°C, washed four times by nutating for 2 minutes in 1×IP buffer, and eluted for 30 minutes at 4°C with 5 μg μL^-1^ 3×FLAG peptide in 1×IP buffer plus 5% digitonin. Note that the concentrations of FLAG peptide and digitonin are higher in our elution step than in the elution step of the original Wang et al protocol. These were included in order to compensate for the higher avidity of membrane fragments bearing multiple FLAG-tagged proteins compared to the individual protein complexes immunoprecipitated in the original protocol, as well as to dissolve the membrane fragments before moving on to protein identification by mass spectrometry.

After elution, samples were denatured by heating at 70°C for 10 minutes, then were loaded into a 4-20% Criterion TGX Precast Mini Protean Gel (BioRad) for electrophoresis at 50 V for 40 min until the protein front moved ∼2.0 cm to separate the FLAG peptides from the eluted proteins. Each lane was cut from below the bottom of the well to just below the bottom of the ladder (<u>∼5 kDa</u>) and gel slices were stored at 4°C until processing for in-gel digestion. Samples were loaded in every other well to decrease cross-contamination between samples. Mass spectrometry was performed as described in (Wang *et al*., 2023). The results of this membrane affinity purification experiment are represented in Fig. **2a** and full results can be found in Supporting Information Dataset **S1**.

**Table 2.** Proteins enriched in at least two membrane affinity purification experiments. Spectral counts in each affinity purification sample for the proteins identified as enriched in at least two out of the three membrane affinity purification experiments. These values are scaled normalized spectral abundance factor values for experiments 1 and 2, which were quantified using label-free quantification; and normalized spectral counts for experiment 3, which was quantified using tandem-mass-tagging. The names of previously-known tubule proteins are bolded.

| Locus | Gene Name | Experiment 1 |  |  |  |  | Experiment 2 |  |  |  |  | Experiment 3 |  |  |  |  |
| --- | --- | --- | --- | --- | --- | --- | --- | --- | --- | --- | --- | --- | --- | --- | --- | --- |
|  |  | RBMP1 | RBMP2 | CPLD9 | CGL129 | fold enrichment | RBMP1 | RBMP2 | CPLD9 | CGL129 | fold enrichment | RBMP1 | RBMP2 | CPLD9 | CGL129 | fold enrichment |
| Cre06.g261750 | **RBMP1** | 5548 | 281 | 0 | 0 | INF | 3678 | 0 | 0 | 0 | INF | 19792 | 1762 | 1159 | 1446 | 8.3 |
| Cre09.g416850 | **RBMP2** | 3669 | 8470 | 0 | 0 | INF | 50 | 727 | 0 | 36 | 21.9 | 3429 | 13989 | 1818 | 1860 | 4.7 |
| Cre02.g143550 | LCI16 | 3312 | 2012 | 0 | 0 | INF | 908 | 312 | 0 | 0 | INF | 2066 | 710 | 478 | 504 | 2.8 |
| Cre09.g415700 | **CAH3** | 5985 | 5454 | 0 | 0 | INF | 1642 | 565 | 549 | 772 | 1.7 | 2955 | 1346 | 702 | 777 | 2.9 |
| Cre06.g259100 | **MITH1** | 704 | 1569 | 0 | 0 | INF | 0 | 399 | 0 | 0 | INF | 1640 | 1370 | 1173 | 1118 | 1.3 |
| Cre10.g452250 | PME1 | 1454 | 4711 | 1196 | 906 | 2.9 | 798 | 1646 | 801 | 563 | 1.8 | 1677 | 2240 | 651 | 555 | 3.2 |

The second affinity purification replicated the first with the following changes: 1) sonication was performed using 6 cycles of (10s at 80% amplitude, 20s rest) on ice; 2) sonicated lysate was clarified at 5,000×g instead of 1,000×g; 3) In place of elution with FLAG peptide and in-gel digestion of proteins, proteins were digested on-bead using S-Trap™ Micro Spin Column Digestion (Protifi); 3) Samples were run on the mass spectrometer for 2 hours instead of 1 hour. The results of this membrane affinity purification experiment are represented in Fig. **2b** and full results can be found in Supporting Information Dataset **S2**.

The third affinity purification was performed like the first with the following changes: 1) sonication was performed using 6 cycles of (10s at 80% amplitude, 20s rest) on ice; 2) sonicated lysate was centrifuged at 5,000×g; 3) after elution, samples were frozen in liquid nitrogen and submitted to the Thermo Fisher Scientific Center for Multiplexed Proteomics at Harvard Medical School for digestion, tandem mass tag (TMT) labeling (Thermo Fisher), and quantitative mass spectrometry (Ting *et al*., 2011; McAlister *et al*., 2012; Li *et al*., 2020). The results of this membrane affinity purification experiment are represented in Fig. **2c** and full results can be found in Supporting Information Dataset **S3**.

### Live-Cell Imaging

For live-cell imaging, strains expressing Venus-tagged proteins were grown at 3% CO_2_ in TP media to a concentration of ∼5×10^5^ cells mL^-1^, then transferred to ambient CO_2_ 16 hours before collecting for imaging at a concentration of ∼1-2×10^6^ cells mL^-1^. 200 µL of each cell culture were placed into the wells of an 8-well µ-slide (Ibidi) and allowed to settle for 5 minutes. The culture media was aspirated to leave a thin layer of cells, and the cells were immobilized for imaging by flowing over 1 mL of 2% low-melting-point TP agarose in each well at ∼40°C, which was allowed to set for 5 minutes at room temperature prior to imaging.

Cells were imaged using a VT-iSIM super-resolution spinning-disk confocal microscope on an Olympus iX83 body equipped with VisiView software, a 60X 1.43 NA objective and a Hamamatsu Orca Quest sCMOS camera in slow mode. Venus fluorescence was excited using a 514 nm laser and collected at 545/50 nm. Chlorophyll autofluorescence was excited using a 642 nm laser and collected at 700/75 nm. Confocal Z-series were acquired with a 300 nm Z-step. The same acquisition settings were used for all strains imaged in this study. Images were imported into Fiji (Schindelin *et al*., 2012) for deconvolution using the Microvolution plugin and final image preparation.

### Transmission Electron Microscopy (TEM)

TEM was performed as described in (He *et al*., 2020) with some modifications. Strains for TEM imaging were grown at 3% CO_2_ in TP media to a concentration of ∼5×10^5^ cells mL^-1^, then transferred to ambient CO_2_ 16 hours before harvesting at a concentration of ∼1-2×10^6^ cells mL^-1^. At least 50×10^6^ cells were harvested by centrifugation at 1,000×g for 5 minutes in 50 mL tubes, resuspended in 1 mL TP, transferred to 1.5 mL screw-top tubes, and centrifuged again for 5 minutes at 1,000 x g. Cells were then resuspended in 2.5% gluteraldehyde in TP media (10 mL 10% gluteraldehyde, 30 mL TP) and nutated at room temperature for 1 hour. After nutation, cells were pelleted at 3,000×g for 1 minute, then washed 3 times by resuspending and nutating in 1 mL ddH_2_O for 5 minutes each, pelleting at 3,000×g after each wash. Samples were then post-fixed and stained with 1 mL freshly-prepared osmium tetroxide solution (1% OsO_4_, 1.5% w/v K_3_[Fe(CN)_6_], 2 mM CaCl_2_) and nutated for 1-2 hours. Tubes were wrapped in aluminum foil during nutation to protect samples from light. Counterstaining with uranyl acetate was found to increase background staining and reduce contrast when looking at membranes, so for all images shown and analyzed here, OsO_4_ was the only stain used. Samples were then serially dehydrated by resuspension and nutation in increasing concentrations of ethanol: 5 minutes in 1 mL each of 30%, 50%, 70%, and 95% ethanol, then 10 minutes in 100% ethanol and 2×10 minutes in 100% acetonitrile. Samples were pelleted for 1 minute at 3,000×g between each resuspension.

After dehydration, samples were embedded in epoxy resin containing 34% Quetol 651, 44% nonenyl succinic anhydride, 20% methyl-5-norbornene-2,3-dicarboxylic anhydride and 2% catalyst dimethylbenzylamine (Electron Microscopy Sciences) over four days, first by embedding overnight in 1:1 acetonitrile:resin (no catalyst) with the screw-top tube open in a fume hood, then by nutating in 1 mL Quetol resin (including catalyst) for four days, spinning down the samples and resuspending in fresh resin once per day. On day 4, the samples were resuspended in 300-500 µL resin, centrifuged at max speed (18,213 x g) for 20 minutes at 30°C in a tabletop centrifuge with a swinging bucket rotor for microfuge tubes, then cured at 60–65°C for 48 hours.

Thin (∼70 nm) sections were prepared from cured resin blocks on a Leica Microtome Ultracut UCT and mounted on carbon film–coated 200 mesh copper TEM grids (Electron Microscopy Sciences) and imaged at the Imaging and Analysis Center, Princeton University, using a CM200 TEM (Philips) or a Talos F200X STEM (ThermoFisher Scientific).

### Growth assays

Cells were grown in liquid TP media at high CO_2_ (∼3%) to a concentration of ∼1-2×10^6^ cells mL^-1^, then harvested by centrifugation by at 1,000×g for 5 minutes. Cultures were resuspended at a density of 6×10^6^ cells mL^-1^, diluted 1:10 and 1:100, and spotted in 5 μL spots onto TP agar. Plates were incubated in 3%, ∼400 ppm, or ∼50 ppm CO_2_ and under 100 or 300 μmol ⋅ m^−2^ ⋅ s^−1^ photons. Plates were imaged after 6 days using a Phenobooth (Singer Instruments).

### Immunofluorescence Staining and Imaging

Immunostaining and imaging were performed as described previously (Yamano & Fukuzawa, 2016). Briefly, cells were centrifuged, washed twice with PBS, and applied to Poly-L-Lysine coated glass slides. Cells were permeabilized with PBS-T and fixed with 4% formaldehyde. Cells were further fixed in 100% ice-cold methanol, except for experiments in which chlorophyll was retained. Cells were rehydrated with PBS, blocked, and stained with α-LCI16 (Yenzym), α-RBMP1 (obtained from the Mackinder Lab; (Adler *et al*., 2024)), or α-FLAG (Cell Signaling 8146T) primary antibodies at a 1:500 dilution. Cells were then washed with PBS-T and stained Goat anti-Rabbit IgG (H+L) Secondary Antibody cross-adsorbed with Alexa Fluor 488 (Invitrogen A-11008) or Alexa Fluor 564 (Invitrogen A-11035), or with F(ab’)2-Goat anti-Mouse IgG (H+L) Secondary Antibody cross-adsorbed to Alexa Fluor 488 (Invitrogen A-11017). Cells were imaged using a VT-iSIM super-resolution spinning-disk confocal microscope on an Olympus iX83 body equipped with VisiView software, a 100x NA 1.35 silicone immersion objective, and a Hamamatsu Orca Quest sCMOS camera in slow mode.

### Whole-cell proteomics

For whole-cell proteomics, strains were inoculated from fresh TAP plates into 15 mL TAP liquid cultures and grown in air at ∼150 μmol ⋅ m^−2^ ⋅ s^−1^ to a concentration of ∼1-2×10^6^ cells mL^-1^. These cultures were used to inoculate 50 mL TP liquid cultures at a density of ∼2.5-5×10^5^ cells mL^-1^ and grown at high CO_2_ (∼3%) at ∼150 μmol ⋅ m^−2^ ⋅ s^−1^ to a concentration of ∼1-2×10^6^ cells mL^-1^. The high CO_2_ cultures were finally used to inoculate 200 mL TP liquid cultures at a density of ∼2.5-5×10^5^ cells mL^-1^, which were grown at low CO_2_ (∼400 ppm) and ∼150 μmol ⋅ m^−2^ ⋅ s^−1^ for 16 hours. Cultures were harvested by centrifugation at 1,000×g and washed three times in PBS. The pellets after the last PBS wash were frozen in liquid nitrogen and stored at −80°C. Samples were then submitted to the Thermo Fisher Scientific Center for Multiplexed Proteomics at Harvard Medical School for digestion, tandem mass tag (TMT) labeling, and quantitative mass spectrometry. Samples were processed as follows: 25 μg of protein from each sample was reduced with TCEP, alkylated with iodoacetamide, and further reduced with DTT. Proteins were precipitated onto SP3 beads to facilitate a buffer exchange into digestion buffer, digested with Lys-C (1:50) overnight at room temperature and with trypsin (1:50) for 6 hours at 37°C, and labelled with TMT 18plex reagents. 2 μL of each sample was pooled and used to shoot a ratio check to confirm complete TMT labelling, and samples were pooled according to the ratios determined by the ratio check. Peptides were desalted using a Sep-pak and fractionated using basic reverse phase HPLC. Six fractions were solubilized, desalted by stage tip, and analyzed on an Orbitrap Fusion Lumos. Peptide spectral matches were filtered to a 1% false discovery rate (FDR) using the target-decoy strategy combined with linear discriminant analysis. The proteins were filtered to a <1% FDR and quantified only from peptides with a summed SN threshold of >180. Comet Search Parameters were as follows: Peptide Mass Tolerance: 50 ppm, Fragment Ion Tolerance: 1.0005, Fragment Bin Offset: 0.4, Theoretical Fragment Ions: 1, Max Internal Cleavage Site: 2, Max differential/Sites: 5, methionine oxidation are used as variable modification. MT Reporter Quant Parameters were as follows: Proteome (MS#) - tolerance = 0.003, ms2 isolation width = 0.7, ms3 isolation width = 1.2. We note that PME1 levels in the *pme1* mutants appear unchanged from wild type; we believe this is due to very low absolute levels of PME1 in wild type preventing accurate quantification. This is supported by the peptide-level quantification, in which PME1 peptide levels were detected at similarly low levels as EPYC1, MITH1, and SAGA1 peptides were detected in their respective mutants, indicating that PME1 peptide levels in both wild-type and the *pme1* mutant strains are low enough that they likely represent noise.

## Results

### Isolation of membrane fragments using affinity purification

The use of affinity purification to identify protein complexes is well-documented (Morris *et al*., 2014; Meyer & Selbach, 2015). Such methods usually involve the use of detergents during cell lysis to dissolve membranes (Behnke & Urner, 2023), such that proteins co-precipitate with the bait protein only if they are part of the same protein complex. This strategy has been used to identify novel protein complexes and elucidate functions for previously uncharacterized genes in *C. reinhardtii* (Mackinder, 2017; Wang *et al*., 2023). However, here we wanted to isolate fragments of tubule membrane to comprehensively identify pyrenoid tubule-localized proteins in an unbiased fashion.

We therefore adapted an existing affinity purification protocol used in *C. reinhardtii* (Mackinder, 2017; Wang *et al*., 2023) to affinity purify intact membrane fragments. Using tagged membrane proteins of known localization (Fig. **1a**) as bait, we performed the affinity purification protocol on cell lysate in which membranes were disrupted by sonication rather than solubilized by detergent (Fig. **1b**). We reasoned that this approach would produce membrane fragments that would allow the identification of proteins that co-localized to the same membrane regions, even if they do not directly interact (Fig. **1c**).

**Figure 1:**
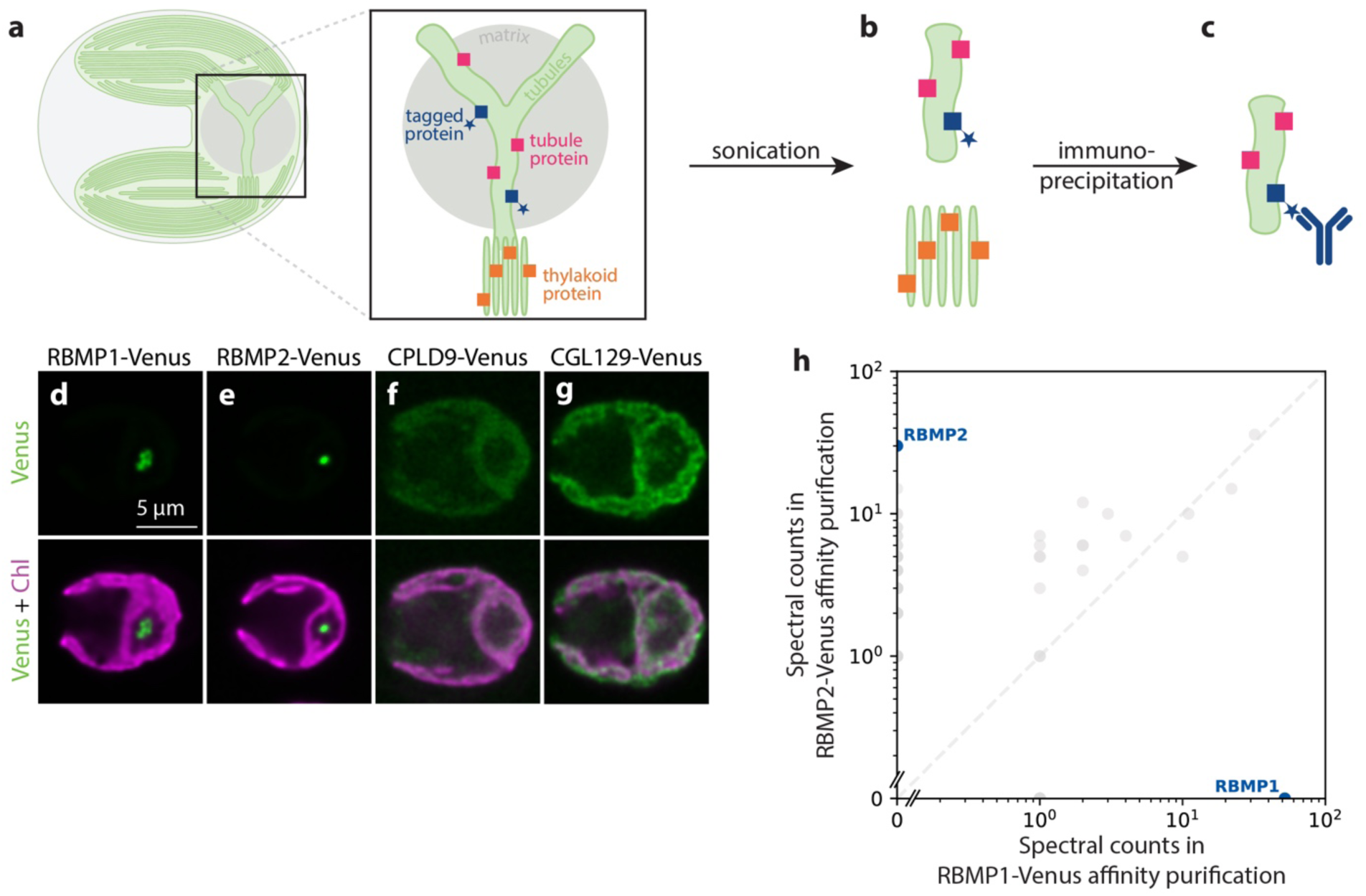
We adapted an affinity purification protocol to isolate intact membranes. a–c. Schematic of the membrane affinity purification protocol. **a.** Cartoon representation of a *C. reinhardtii* cell, with the pyrenoid in the base of the cup-shaped chloroplast. Inset: The pyrenoid is traversed by tubule membranes which are contiguous with—but structurally distinct from—the photosynthetic thylakoids. Colored squares represent the proteins localized to the membranes: blue squares with a star represent known tubule proteins tagged with a FLAG tag, pink squares represent all other proteins localized to the tubules, and orange squares represent proteins localized to the thylakoid membranes. **b.** By disrupting cellular membranes using sonication, we created small, intact fragments of pyrenoid tubules (top) and thylakoids (bottom). **c.** Membrane fragments containing tagged proteins were enriched by immunoprecipitation using an anti-FLAG antibody, and their protein composition was determined by mass spectrometry. **d–g.** Confocal fluorescence images showing the localization of the FLAG-tagged proteins used as baits for the membrane affinity purification. Chlorophyll, a proxy for the presence of thylakoid membranes, is shown in magenta. The pyrenoid is visible as a “hole” in the chlorophyll fluorescence at the base of each cell (on the right side of each image). Venus signal is shown in green, representing the localization of the tubule **(d–e)** or thylakoid **(f–g)** bait proteins. **h.** Results of a detergent-based affinity purification show that the tubule bait proteins RBMP1-Venus and RBMP2-Venus do not co-precipitate using traditional affinity purification methods.

We selected four bait proteins on which to perform this protocol: RBMP1-Venus-3×FLAG and RBMP2-Venus-3×FLAG (Fig. **1d,e**), which localize to the pyrenoid tubules (Meyer *et al*., 2020b); and CPLD9-Venus-3×FLAG and CGL129-Venus-3×FLAG (Fig. **1f,g**), which localize to the thylakoid membranes and are excluded from the tubules (Wang *et al*., 2023). RBMP1 and RBMP2 localize to distinct regions of the tubules—RBMP1 to the tubule periphery and RBMP2 to the central reticulated region (Meyer *et al*., 2020b, Supporting Information Fig. **S1**)—thus, we reasoned that together they should provide coverage for the whole tubule network. As expected from their distinct localizations, RBMP1 and RBMP2 did not co-precipitate via traditional detergent-based affinity purification (Fig. **1h**, Supporting Information Dataset **S4**).

### A membrane affinity purification–based tubule proteome identifies known and novel proteins

We performed three iterations of the membrane affinity purification experiment, each with slight modifications to the protocol to try and improve the enrichment or identification of membrane proteins. Samples from the first two experiments were quantified using label-free quantification (LFQ), identifying 58 and 135 proteins, respectively, that were at least 2-fold enriched in the tubule samples relative to the control samples (Fig. **2a-b**, Supporting Information Datasets **S1– 2**). Quantification using LFQ, however, can suffer from Poisson noise for lower-abundance proteins, leading some low-abundance proteins to appear enriched or depleted in a given sample due to differences in random run-to-run peptide sampling rather than differences in actual abundance in each sample (O’Connell *et al*., 2018). We therefore set a minimum abundance threshold for proteins to be considered hits in those experiments, which reduced the number of 2-fold-enriched hits in the first two experiments to 22 and 92, respectively. To further address the problem of noisy results for low-abundance proteins, the third iteration of the membrane purification experiment used tandem mass tag (TMT) labeling instead of LFQ. TMT labelling allows all samples to be multiplexed and run simultaneously, so that any random sampling of low-abundance peptides applies to all samples equally, leading to more accurate quantification of lower-abundance proteins and obviating the need for a minimum abundance threshold. This reduction in noise led to fewer hits in the TMT experiment, which only had 14 proteins that were 2-fold enriched in the tubule samples (Fig. **2c**, Supporting Information Dataset **S3**).

**Figure 2:**
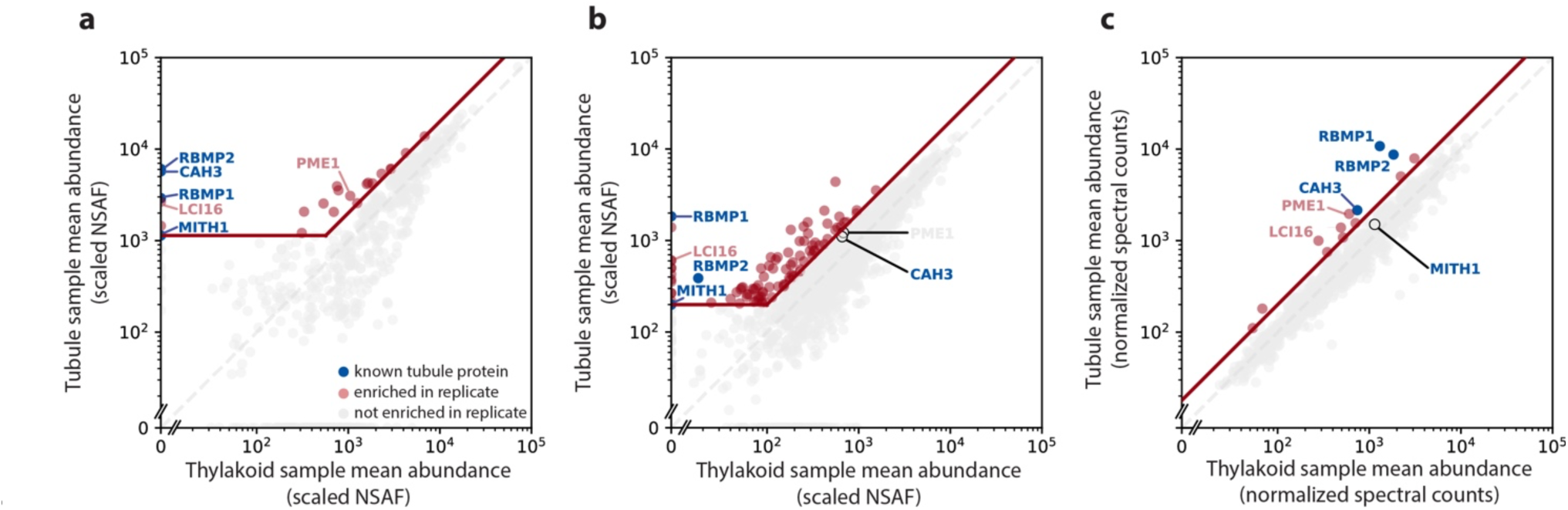
The membrane affinity purification experiment identified both known and novel tubule proteins. Protein abundances in tubule samples vs. thylakoid samples for the three membrane affinity purification experiments. Blue points represent the known tubule proteins RBMP1, RBMP2, CAH3, and MITH1. The grey dashed lines represent y=x, at which proteins are equally abundant in both the tubule and thylakoid samples. The diagonal red line represents 2-fold enrichment in the tubule samples. The first two experiments **(a–b)** were quantified using label-free quantification, leading to significant noise among lower-abundance proteins. For these experiments, hits (red points) were defined as exceeding 2-fold enrichment in the tubule sample and exceeding a lower abundance threshold (horizontal red line). Proteins in the third membrane affinity purification **(c)** were quantified using tandem mass tagging, leading to better quantification of low-abundance proteins. Thus, only 2-fold enrichment was required to be considered a hit. Proteins that were hits in at least two experiments can be found in Table **2**.

We consolidated the results of all three experiments into one list of high-confidence tubule proteins by selecting only the proteins that were at least 2-fold tubule-enriched in at least two out of three of the membrane purifications. There were only six such proteins, which are listed in Table **2**. Four of these six proteins are previously-known tubule proteins: the two bait proteins RBMP1 and RBMP2, the carbonic anhydrase CAH3 (Sinetova *et al*., 2012), and MITH1 (Hennacy *et al*., 2024).

While these four proteins do not represent all previously-known tubule-localized proteins, the nature of our experiment is such that we measured protein enrichment relative to the rest of the thylakoid membranes, so our method is less sensitive to proteins that localize to both the tubules and thylakoids. Many tubule-enriched proteins, such as PSAH, TEF14, and HCF136, are observed in the tubules but are also present in the thylakoids (Mackinder, 2017; Wang *et al*., 2023), while others, like CAS1, re-localize from throughout the chloroplast to the tubules in a condition-dependent manner (Wang *et al*., 2016). For such proteins, we expect that some protein may still be present in the thylakoids under our experimental conditions, decreasing their tubule enrichment factors. Thus, Table **2** does not represent all proteins present in the tubules, only those which are most highly enriched relative to their abundance in the thylakoids in our assay.

Importantly, our results are consistent with the enrichment of proteins based on membrane proximity rather than direct protein-protein interaction, as we were able to identify proteins not seen in traditional detergent-based affinity purification. Neither MITH1 nor CAH3 co-immunoprecipitated with RBMP1 or RBMP2 in a detergent-based affinity purification (Supporting Information Dataset **S4**), yet both are enriched in our tubule samples (Fig. **2**, Table **2**). The presence of CAH3, a pyrenoid tubule-enriched luminal protein with no predicted transmembrane domains (Sinetova *et al*., 2012), also suggests that we are able to capture proteins within the membrane lumen. Furthermore, RBMP1 and RBMP2 did not co-precipitate in detergent-based affinity purification (Fig. **1h**, Supporting Information Dataset **S4**), whereas they did co-precipitate in our first and third membrane purifications (Table **2**).

Taken together, these results indicate that our protocol is able to isolate and identify tubule membrane-enriched proteins that are not captured by traditional affinity purification approaches. These observations gave us confidence in the likelihood that the other two proteins in our top hits were also tubule-localized. These proteins were Low-CO_2_-induced 16, also known as Early Light-Induced 4 (LCI16/ELI4; Cre02.g143550) and Cre10.g452250, which we named Pyrenoid Membrane–Enriched 1 (PME1). Given these proteins’ presence in our tubule proteome and their relative lack of characterization, we sought to validate their tubule localization and determine if they impact pyrenoid tubule biogenesis.

### Fluorescently-tagged LCI16 localizes to the pyrenoid-traversing membranes

We created constructs to express each novel tubule candidate gene tagged with a Venus fluorophore and transformed each construct into wild-type cells (CMJ030). Multiple PME1-Venus-3×FLAG constructs, using both the native promoter and the highly-expressing PSAD promoter, consistently failed to yield detectable levels of Venus fluorescence, even among antibiotic-resistant colonies that appeared to have been successfully transformed. Antibodies raised against PME1 also failed to produce a PME1-specific signal, preventing us from using them effectively to localize PME1. These challenges may be related to PME1’s low native abundance (Fig. **6**, Supporting Information Fig. **S2a**). Thus, we focused our efforts on characterizing LCI16-Venus-3×FLAG transformants, which yielded detectable fluorescence.

We observed via fluorescence microscopy that LCI16-Venus-3×FLAG localizes to the center of the pyrenoid (Fig. **3a**). Its localization pattern was concentrated near the center of the pyrenoid, more similar to the pattern of known tubule proteins RBMP1 and RBMP2 (Fig. **1d,e**) than to the pattern of matrix-localized proteins such as Rubisco or EPYC1 (Meyer *et al*., 2020b). We attempted to determine the localization of LCI16 more precisely by co-localizing it with RBMP2-Venus using an antibody raised against LCI16. However, the α-LCI16 antibody stained the entire chloroplast, including the pyrenoid (Supporting Information Fig. **S3b**). This could indicate that LCI16 localizes to the entire chloroplast, or that the antibody is not specific to LCI16. To distinguish between these possibilities, we stained a mutant lacking LCI16 expression with the α-LCI16 antibody and compared it to α-LCI16 staining in wild-type cells. In the *lci16* mutant strain, the non-pyrenoidal signal remained, but the pyrenoid signal was no longer observed (Supporting Information Fig. **S3e**), indicating that the non-pyrenoidal signal in wild-type cells was likely non-specific. Futhermore, the loss of α-LCI16 pyrenoid staining in the *lci16* mutant is consistent with a pyrenoid localization for LCI16, as observed in the LCI16-Venus-3×FLAG strain. Based on LCI16’s enrichment in our tubule membrane proteome and the localization pattern observed for LCI16-Venus-3×FLAG, we conclude that LCI16 localizes to pyrenoid-traversing membranes.

**Figure 3:**
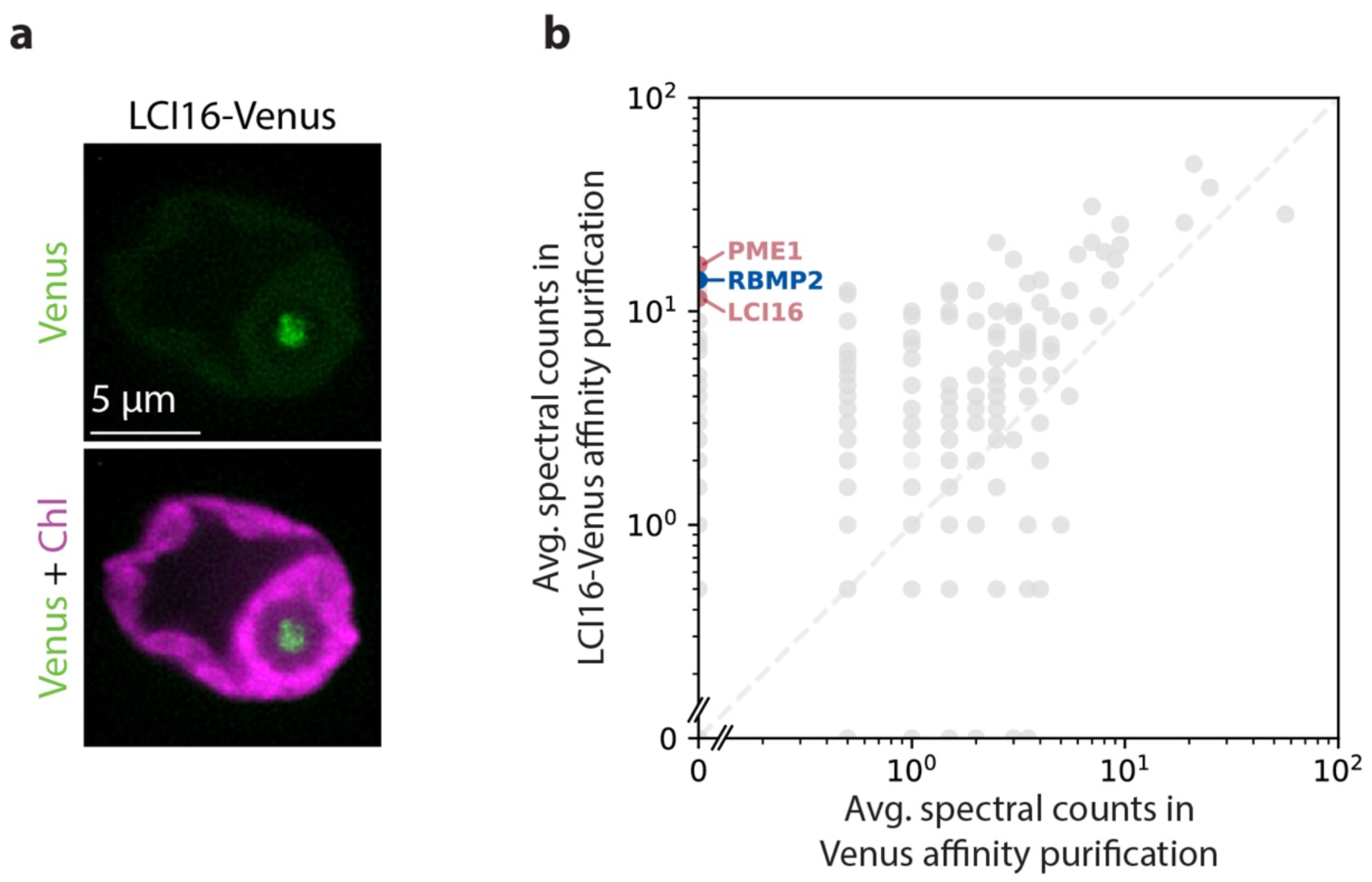
LCI16-Venus localizes to the pyrenoid tubules and co-precipitates RBMP2 and PME1. **a**. Confocal images showing the localization of LCI16-Venus in the pyrenoid tubules. Green represents LCI16-Venus signal and magenta represents chlorophyll. **b**. Average spectral counts for a detergent-based affinity purification of LCI16-Venus compared to a control affinity purification of Venus alone. The blue point represents the known tubule protein RBMP2, while the red points represent the novel tubule components LCI16 and PME1.

### LCI16 physically interacts with RBMP2 and PME1

To further investigate the role of LCI16 in the tubule membranes, we performed a traditional detergent-based affinity purification using LCI16-Venus-3×FLAG as bait to identify its protein interactors (Fig. **3b**, Supporting Information Dataset **S5**). In this experiment, the top two interactors of LCI16 were PME1 and RBMP2. This indicates that these three proteins physically interact, directly or indirectly, providing additional support for PME1’s tubule localization.

Our previous detergent-based RBMP2-Venus-3×FLAG affinity purification did not identify LCI16 or PME1 as highly enriched (Supporting Information Dataset **S4**). However, previously-published proteomic data from diurnally-grown cells (Strenkert *et al*., 2019) suggest that RBMP2 is more abundant than LCI16 or PME1 (Supporting Information Fig. **S2**), raising the possibility that a large proportion of LCI16 and PME1 may be bound to RBMP2 even while the majority of RBMP2 is not bound to LCI16 or PME1.

### Transmission electron microscopy (TEM) of insertional mutants in LCI16 and PME1 indicates that neither protein is required for normal tubule morphology

To assess whether LCI16 and PME1 play structural roles in pyrenoid tubule biogenesis, we examined pyrenoid morphology in mutant strains lacking expression of each protein using transmission electron microscopy (TEM). We obtained three mutant alleles for both LCI16 and PME1, with four mutants coming from the CLiP collection (Li *et al*., 2019) and two mutants coming from the more recently-developed CLiP2 collection (Lunardon *et al*., 2024). We confirmed the insertion sites in all six mutants by PCR (Supporting Information Fig. **S4**) and examined all mutants by electron microscopy.

Unlike *mith1* and *saga1* mutants, which have the most severe tubule-related defects observed in *C. reinhardtii* to date—with most Rubisco condensates entirely lacking tubules (Hennacy *et al*., 2024)—the *lci16* and *pme1* mutants examined by TEM showed wild-type pyrenoid morphology. We regularly observed only one pyrenoid per cell, and we could observe central reticulated regions (Fig. **4b,e,h**) and peripheral cylindrical tubules with minitubules (Fig. **4c,f,i**, Supporting Information Fig. **S5**) that were indistinguishable from those in wild-type cells. These results indicate that LCI16 and PME1 are not required for normal tubule morphology under our culture conditions and suggest that they function in processes unrelated to tubule biogenesis.

**Figure 4:**
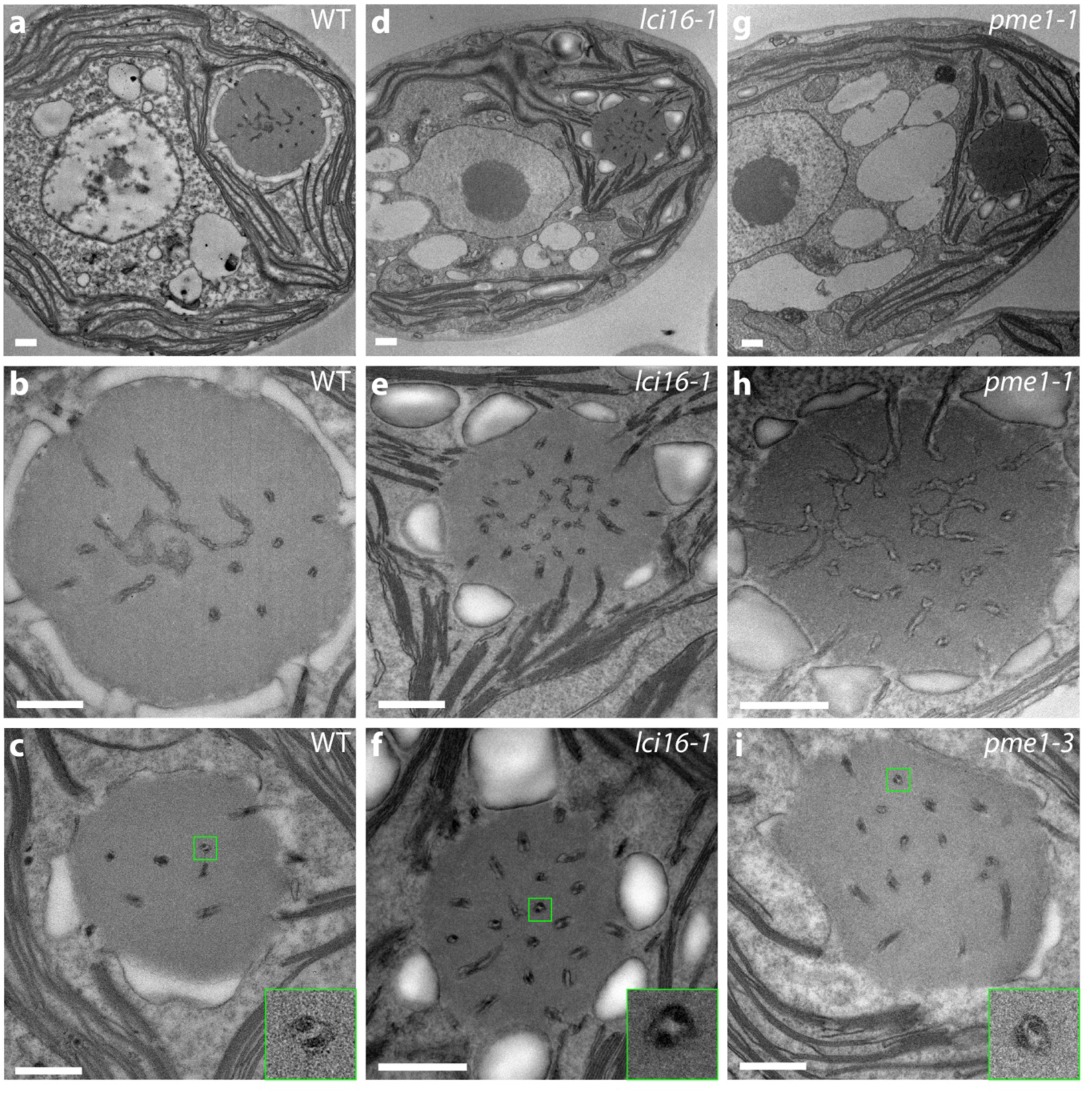
Transmission electron microscopy indicates normal pyrenoid morphology in insertional mutants of *lci16* and *pme1*. Electron micrographs of wild type **(a–c)**, *lci16* **(d–f)**, and *pme1* **(g–i)** cells. Images of central slices of whole cells **(a,d,g)** show the single pyrenoid in its stereotyped location at the base of the cup-shaped chloroplast. Higher-magnification images of the pyrenoids **(b–c, e–f, h–i)** show tubules in all of the pyrenoids, including both the central reticulated region, characterized by three-way membrane junctions **(b,e,h)**, and the peripheral, minitubule-containing tubules **(c,f,i)**. 4× insets in **(c,f,i)** show individual tubules with visible minitubules. Pyrenoids in **(b,e,h)** represent the same cells as shown in **(a,d,g)**. Scale bars 500 nm.

### *lci16* and *pme1* mutants have normal growth under CCM-requiring conditions

We further investigated the role of LCI16 and PME1 in CCM function by performing spot tests on the *lci16* and *pme1* insertional mutants under increasingly CO_2_-limiting growth conditions (Fig. **5**). We compared the growth of mutants to the growth of their respective wild-type background strains, as well as to the growth of *saga1* and *mith1* mutants, which are known to have CCM defects (Itakura *et al*., 2019; Hennacy *et al*., 2024).

**Figure 5:**
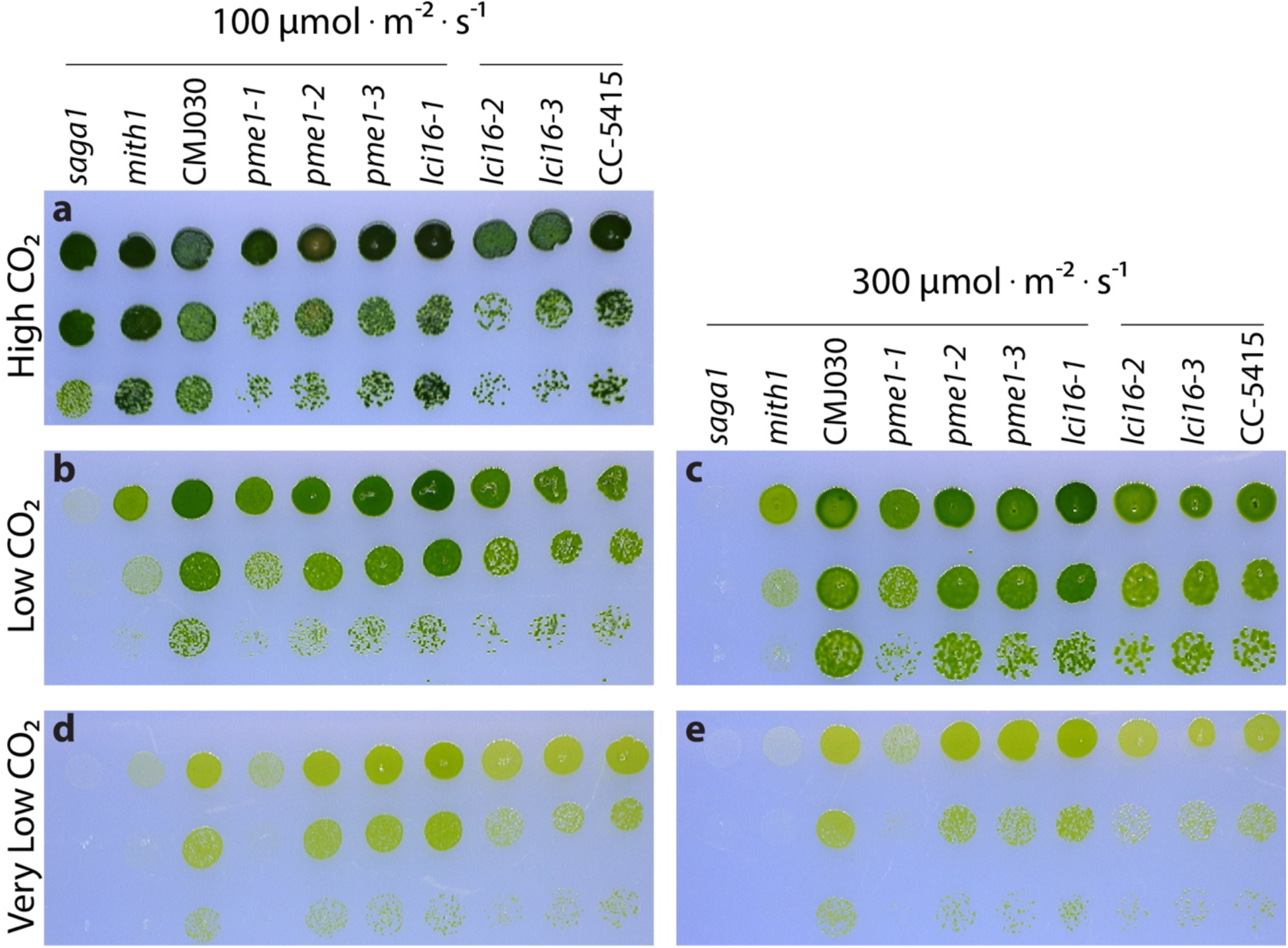
LCI16 and PME1 are not required for normal growth under CCM-inducing conditions. Spot tests of wild-type strains, *lci16* and *pme1* mutants, and known CCM mutants *saga1* and *mith1* under high (3%) **(a)**, low (∼400 ppm) **(b-c)**, and very low (∼40 ppm) **(d-e)** CO_2_ conditions under 100 **(a,b,d)** and 300 **(c,e)** μmol ⋅ m^−2^ ⋅ s^−1^ of photons. All mutant strains are in the CMJ030 wild-type background strain except for *lci16-2* and *lci16-3*, which are in the CC-5415 background. Horizontal lines over the genotypes indicate strains that share the same wild-type background.

All strains grew as well as wild-type under high CO_2_ (Fig. **5a**). As expected, the *saga1* and *mith1* mutants suffered increasingly severe growth defects as the CO_2_ level decreased and the light intensity increased (Fig. **5b-e**). However, the *lci16* and *pme1* mutants grew as well as their respective wild-type strains under all conditions tested, with the exception of one *pme1* mutant, *pme1-1*, which had a mild growth defect under very low CO_2_. Given that two of the three *pme1* mutants displayed no CCM growth defect, it is most likely that a second, off-site mutation in *pme1-1* is responsible for its growth defect. Together, these results suggest that neither LCI16 nor PME1 is required for proper CCM function under the conditions tested.

### Proteomes of the *lci16* and *pme1* mutants are similar to those of known CCM mutants

Since the *lci16* and *pme1* mutants had no defect in their tubules and no visible growth defect under the conditions tested, we sought to determine whether they impact the proteome. We used mass spectrometry to determine whole-cell proteomes of two alleles each of *lci16* and *pme1*; both of their wild-type background strains; and four mutants with known CCM defects— *cia5*, *epyc1*, *mith1*, and *saga1* (Fig. **6**, Supporting Information Datasets **S6-7**). Many known CCM-related proteins showed similar patterns of regulation across all the known CCM mutants—the putative bicarbonate transporter BST2 was upregulated, while the bicarbonate transporters HLA3 and LCIA and the carbonic anhydrases CAH1, CAH2, CAH5, and LCID were downregulated.

**Figure 6:**
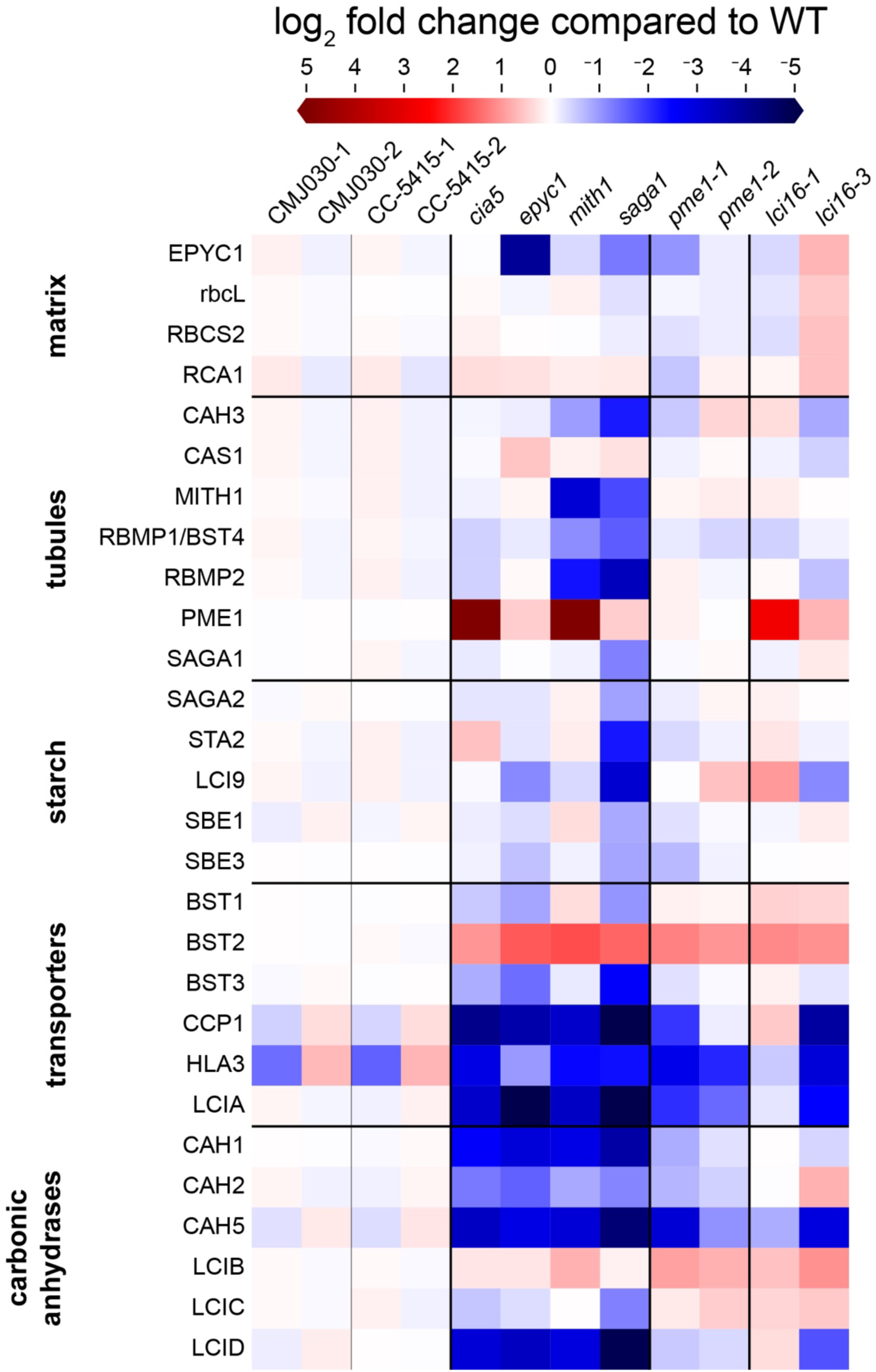
Comparison of CCM-related protein abundance in known CCM mutants and *lci16* and *pme1* mutants. Mass spectrometry was used to determine the whole-cell proteome of the known CCM mutants *cia5*, *epyc1*, *mith1*, and *saga1*; two alleles each of *lci16* and *pme1*; and two biological replicates of each strain’s wild-type background (all strains are in the CMJ030 background except for *lci16-3*, which is in the CC-5415 background). All strains were pre-cultured in TP high CO_2_ (3% CO_2_), then diluted and shifted to low CO_2_ levels (∼400 ppm CO_2_) for 16 hours prior to harvesting for mass spectrometry. The heatmap shows the relative abundance of CCM-related proteins (rows) in each genotype (columns), expressed as the log_2_ fold-change of each protein relative to its wild-type parental strain (averaged between both biological replicates). Proteins are grouped into localization- and function-based groups, denoted by horizontal black lines. Proteins that were quantified based on only one peptide are excluded from the heatmap, but can be found, along with the full dataset, in Supporting Information Datasets **S6-7**.

The *pme1* and *lci16* mutants had similar proteomic profiles to the known CCM mutants, exhibiting upregulation of BST2 and downregulation of HLA3, LCIA, and CAH3. This similarity in proteomic profile is not due to a general growth defect, as the proteomic changes were observed in alleles of *pme1* and *lci16* that have wild-type-like growth under all conditions tested. Considering that disrupting genes in the same pathway typically leads to similar perturbations to the proteome (Kafri *et al*., 2023), we interpret the similarity of *pme1* and *lci16* proteomes to those of known CCM mutants as an indication that PME1 and LCI16 share a function with CIA5, EPYC1, MITH1, and SAGA1 as perceived by cellular regulatory networks.

## Discussion

### Benefits and limitations of the membrane affinity purification protocol

In this study, we enriched pyrenoid-traversing membrane fragments from *C. reinhardtii* and identified their associated proteins. From this dataset, we further characterized two novel tubule protein candidates, LCI16 and PME1.

Our membrane affinity purification approach overcame the challenge that pyrenoid tubules are contiguous with—and vastly less abundant than—the thylakoid membranes, impeding their isolation by traditional membrane fractionation methods (Bailyes *et al*., 1997; Avila *et al*., 2015). Our protocol overcame this challenge by enriching specific membrane subdomains based on the presence of a known resident protein, allowing us to identify novel protein components of the chosen membrane starting with only the identity of two of its protein components. The limitations of our implementation included low-number sampling noise due to the label-free quantification used in the first two experiments, which limited our ability to detect lower-abundance proteins, and changes to the protocol over the three iterations, which we think contributed to variability in hits between the experiments. With further improvements, we believe our approach could be applied to determine the protein composition of other membrane domains on a sub-organellar level.

### Possible functions for the novel tubule components LCI16 and PME1

One of the proteins identified by our protocol is Low-CO_2_-Induced 16 (LCI16), also known as Early Light-Induced 4 (ELI4). LCI16 has a light-harvesting complex (LHC)-like domain homologous to those found in the stress-related LHCs (LHCSR) and Early Light-Induced Protein (ELIP) families, which are characterized by a 3-helix chlorophyll-binding domain (Rochaix & Bassi, 2019; Levin & Schuster, 2023). These proteins are present across plant and algal species; are often induced by dark-to-light transitions, high light, and/or other abiotic stresses; and typically function in photoprotection via non-photochemical quenching (NPQ), as well as assembly of new photosystems or repair of damaged photosystems (Montané & Kloppstech, 2000; Rochaix & Bassi, 2019; Levin & Schuster, 2023). Unlike the ELIPs, however, LCI16 retains relatively high mRNA levels throughout the night, and its expression profile is much more similar to that of RBMP2 than any of the other ELIPs (Supporting Information Fig. **S2**). Given its transcriptional similarity to and physical interaction with RBMP2, it is likely that LCI16 plays a more pyrenoid-specific role than other ELIP proteins. Further, the fact that LCI16 is not required for growth in the light indicates that any light energy absorbed by chlorophyll in the tubules must be used or dissipated by proteins other than LCI16. These could be other, redundant light stress proteins, or they could be tubule-localized photosystem components (Mackinder, 2017; Wang *et al*., 2023), which may pump protons into the tubule lumen to drive the conversion of HCO_3_^-^ to CO_2_ by CAH3. Further work on LCI16 and other tubule-localized light-responsive proteins will be required to determine if the tubules perform any such light-dependent functions beyond serving as the interface between the Rubisco matrix and the conversion of CO_2_ by CAH3.

The other novel tubule protein we identified and characterized, PME1, co-precipitates with LCI16 (Fig. **3b**). While we could not identify any homologs of PME1, it has several non-canonical PRICHEXTENSN-like motifs reminiscent of similar proline-rich domains previously described in proteins associated with plant cell wall stability (Showalter *et al*., 2010; Goodstein *et al*., 2012; Guo *et al*., 2014; Craig *et al*., 2023). The C-terminus of PME1 is predicted by Alphafold 3 (Abramson *et al*., 2024) to fold into a series of beta sheets followed by a long amphipathic helix which may oligomerize as a coiled coil homo-trimer (Supporting Information Fig. **S6a-b**). Amphipathic helices and coiled coils are common oligomerization domains (Truebestein & Leonard, 2016), but they are also common among several families of membrane-binding and -shaping proteins, such as BAR domain proteins (Gallop *et al*., 2006; Jao *et al*., 2010) and reticulons (Brooks *et al*., 2021), suggesting a potential mechanism for membrane binding in the C-terminal domain of PME1.

Our whole-cell proteome experiments also found that PME1 is highly upregulated in the *cia5* and *mith1* mutants (Fig. **6**). This is unusual considering that most CCM-related proteins are downregulated in these mutants—and because under most conditions, PME1 is expressed at very low, often undetectable, levels (Strenkert *et al*., 2019; Kafri *et al*., 2023). These observations raise the possibility that PME1 contributes to CCM regulation, e.g. by promoting expression of CCM-related transporters and carbonic anhydrases in response to inadequate CCM function. Further experiments will be required to determine if PME1 plays such a regulatory role, and under what conditions, if any, it is required for full CCM function.

Taken together, our results identify LCI16 and PME1 as novel pyrenoid-traversing membrane components, initiate their characterization, and suggest directions for future research into their functions. Our pyrenoid-traversing membrane proteome identifies known components and additional candidates, helping to prioritize proteins for future study. We hope these findings will ultimately contribute toward a mechanistic understanding of the pyrenoid, an organelle with a central role in the global carbon cycle.

## Acknowledgments

We would like to thank Saw Kyin and Henry Shwe of the Princeton University Proteomics Core and the Thermo Fisher Scientific Center for Multiplexed Proteomics at Harvard Medical School (https://tcmp.hms.harvard.edu) for advice and help performing mass spectroscopy; John Schreiber and Paul Shao of the Princeton Imaging & Analysis Center for advice and technical support for electron microscopy; Michelle Warren-Williams for media preparation and propagating strains; Victoria L. Crans and Micah Burton for sharing reagents, experimental assistance, and feedback on the manuscript; Ned Wingreen, Fred Hughson, and members of the Jonikas laboratory for project feedback and helpful conversations; and Marie Bao, as part of Life Science Editors, for help with editing the manuscript. The authors acknowledge the use of the Imaging and Analysis Center (IAC) operated by the Princeton Materials Institute at Princeton University, which is supported in part by the Princeton Center for Complex Materials (PCCM), a National Science Foundation (NSF) Materials Research Science and Engineering Center (MRSEC; DMR-2011750). Research reported in this paper was supported by the Howard Hughes Medical Institute, Bill and Melinda Gates Foundation and United Kingdom Foreign, Commonwealth & Development Office grant INV-054558, National Science Foundation grants MCB-2410354 and MCB-1914989, and NIGMS of the National Institutes of Health grants 1R01GM140032-01 and T32GM007388. The content is solely the responsibility of the authors and does not necessarily represent the official views of the National Institutes of Health.

## Author contributions

EF and MCJ conceived of and planned the project. EF, ERC, KD, and SLE performed the membrane purification experiments and EF analyzed the mass spectrometry data. EF performed the TEM imaging, growth assays, and sample preparation for whole cell proteomics. LW performed detergent-based affinity purifications; LW and AG performed fluorescence microscopy. AL, WP, and CP generated and identified the CLiP2 mutants. EF and MCJ wrote the manuscript.

## Data Availability

The data that supports the findings of this study are available in Supporting Information Datasets **S1-7** included with this article.

---

**Supporting Information Figure S1:**
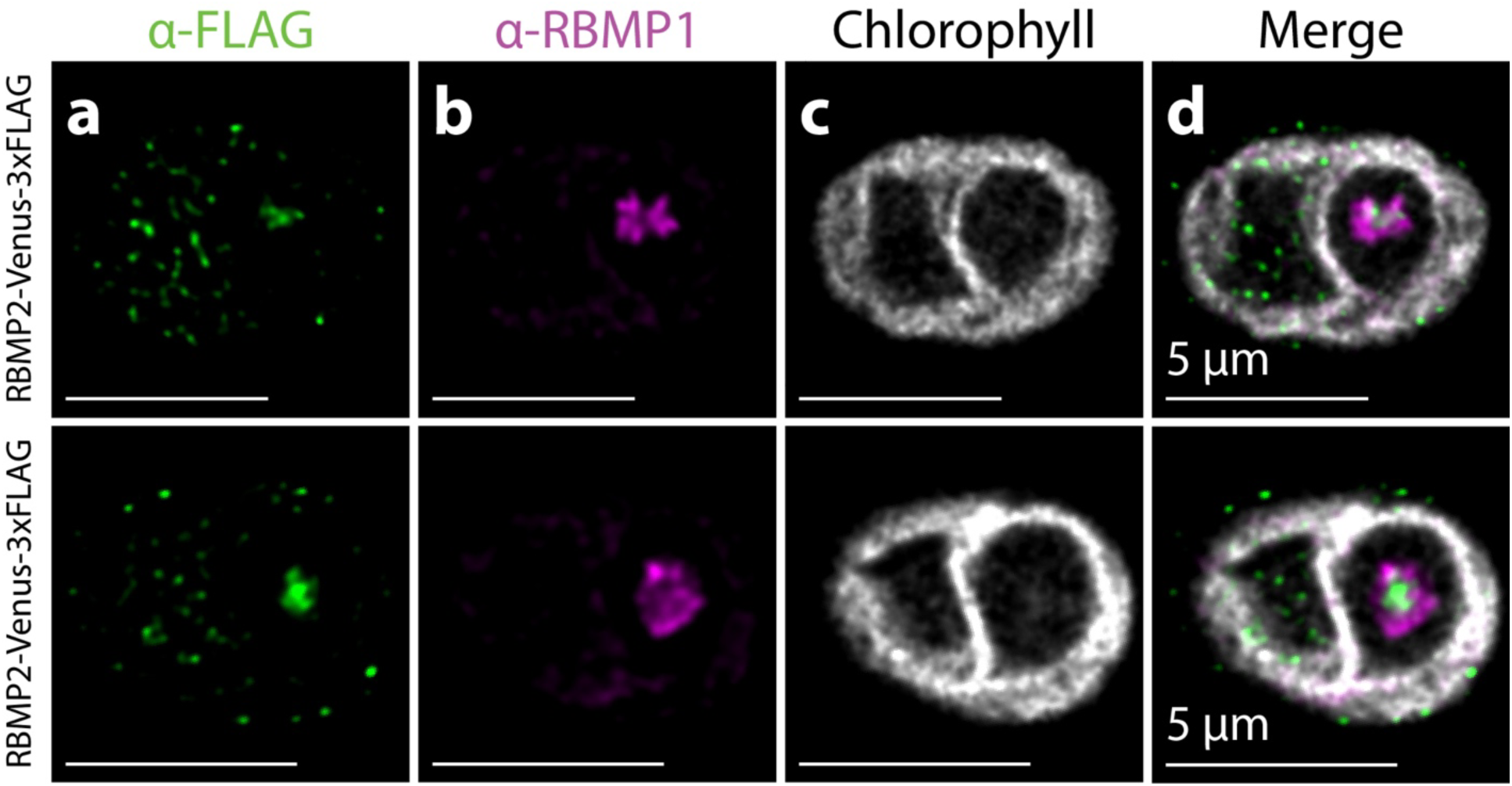
RBMP1 and RBMP2 localize to different regions of the pyrenoid tubules. Wild-type cells expressing RBMP2-Venus-3×FLAG were immunostained using α-FLAG **(a)** and α-RBMP1 **(b)** antibodies. Each row represents an independent cell from the same RBMP2-Venus-3×FLAG strain. A gap in the chlorophyll signal **(c)** in the base of each cell (on the right side of each image) represents the location of the pyrenoid. Merging the channels **(d)** shows the distinct localization of RBMP1 and RBMP2.

**Supporting Information Figure S2:**
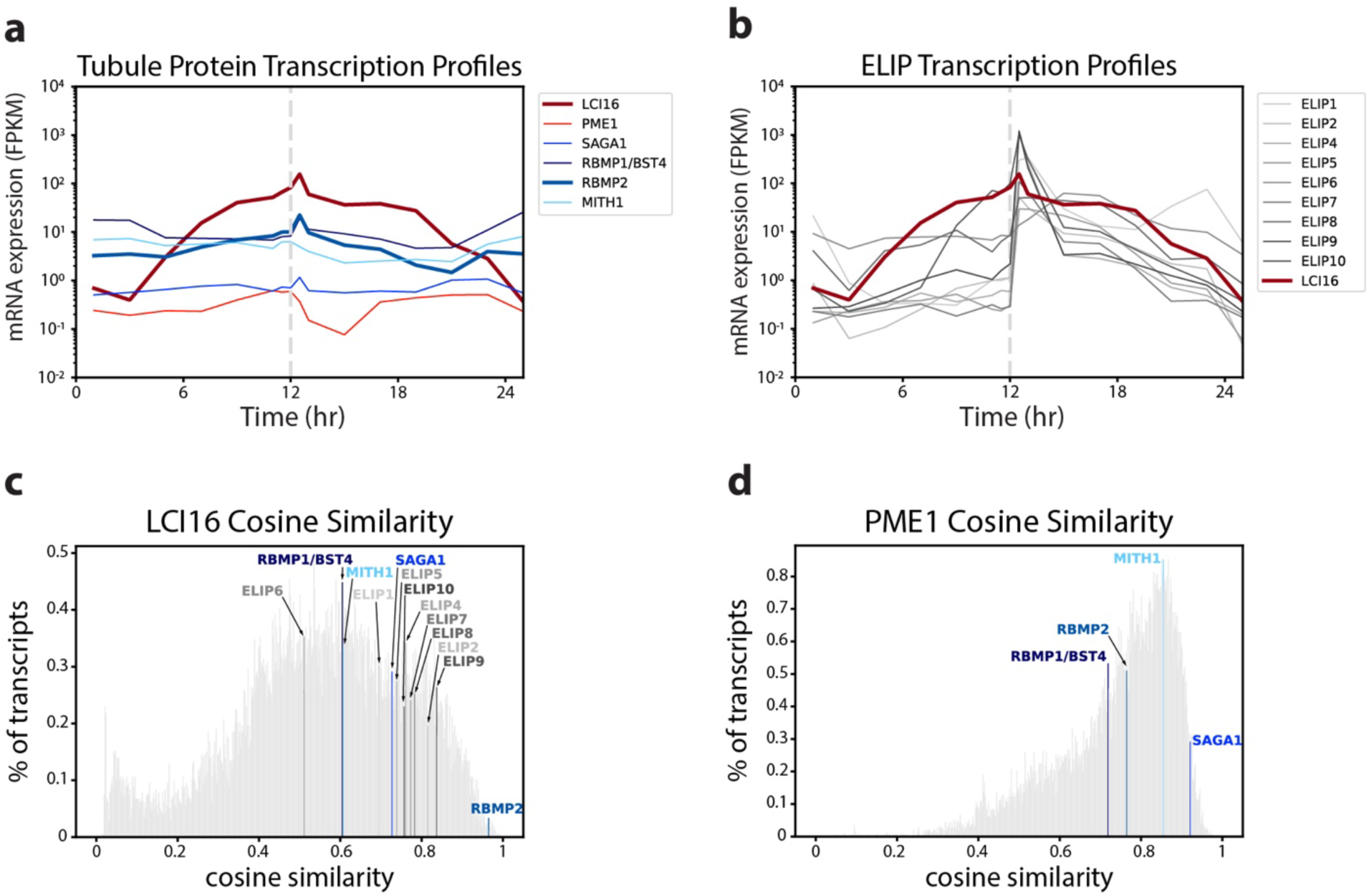
Comparison of LCI16’s and PME1’s expression profiles in diurnally-grown cells to the expression profiles of known tubule proteins and ELIP proteins. Plots showing transcriptomic and proteomic data from diurnally-grown *C. reinhardtii* cells measuring the abundance of mRNA (**a–b**) levels (Strenkert *et al*., 2019) of known pyrenoid tubule proteins (blue lines; **a**) and the Early Light-Induced Proteins (ELIPs, grey lines; **b**) with which LCI16 shares homology. The transcription profile of LCI16 is shown in dark red in both panels. Hours 0 and 24 correspond to the onset of dark and hour 12 corresponds to the onset of light. (**c–d)** Histograms showing the cosine similarity of the Strenkert transcriptome profiles to the LCI16 (**c**) and PME1 (**d**) transcriptome profiles. The cosine similarity calculation considers each gene’s transcription profile as a 16-dimensional vector (each dimension corresponding to one of the 16 timepoints measured) and measures the similarity between transcription profiles by calculating the cosine of the angle between the profiles’ 16-D vectors. A cosine similarity of 1 corresponds to vectors that are parallel, i.e. identical transcriptome profiles. Note that because the angle between vectors is dependent on their direction, rather than their magnitude, this metric measures the similarity of the shape of the transcription profiles (i.e., which timepoints are up- and down-regulated), rather than the absolute transcript abundance at each timepoint. of Bins containing proteins known tubule proteins or ELIPs are colored as in panels **a–b**.

**Supporting Information Figure S3:**
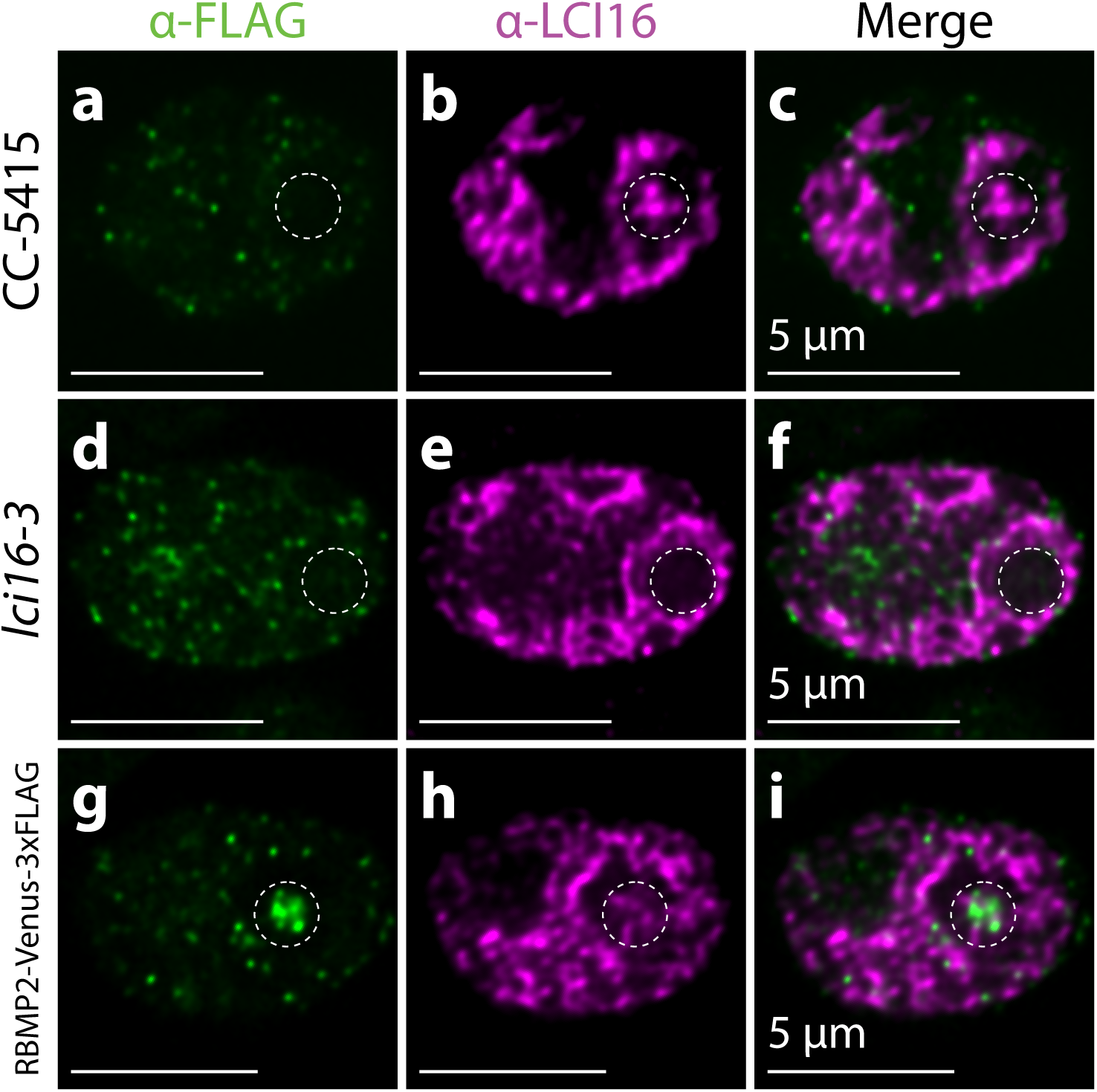
LCI16 antibody staining is non-specific but consistent with LCI16 tubule localization. α-FLAG and α-LCI16 immunostaining in wild-type **(a-c)**, *lci16-3* **(d-f)**, and RBMP2-Venus-3xFLAG **(g-i)** cells. Pyrenoids are denoted with dashed circles. The α-LCI16 antibody stains the entire chloroplast, including the pyrenoid, in wild-type **(b)** and RBMP2-Venus-3xFLAG cells **(h)**. However, in the *lci16-3* mutant, the α-LCI16 signal is notably absent from the pyrenoid **(e)**.

**Supporting Information Figure S4:**
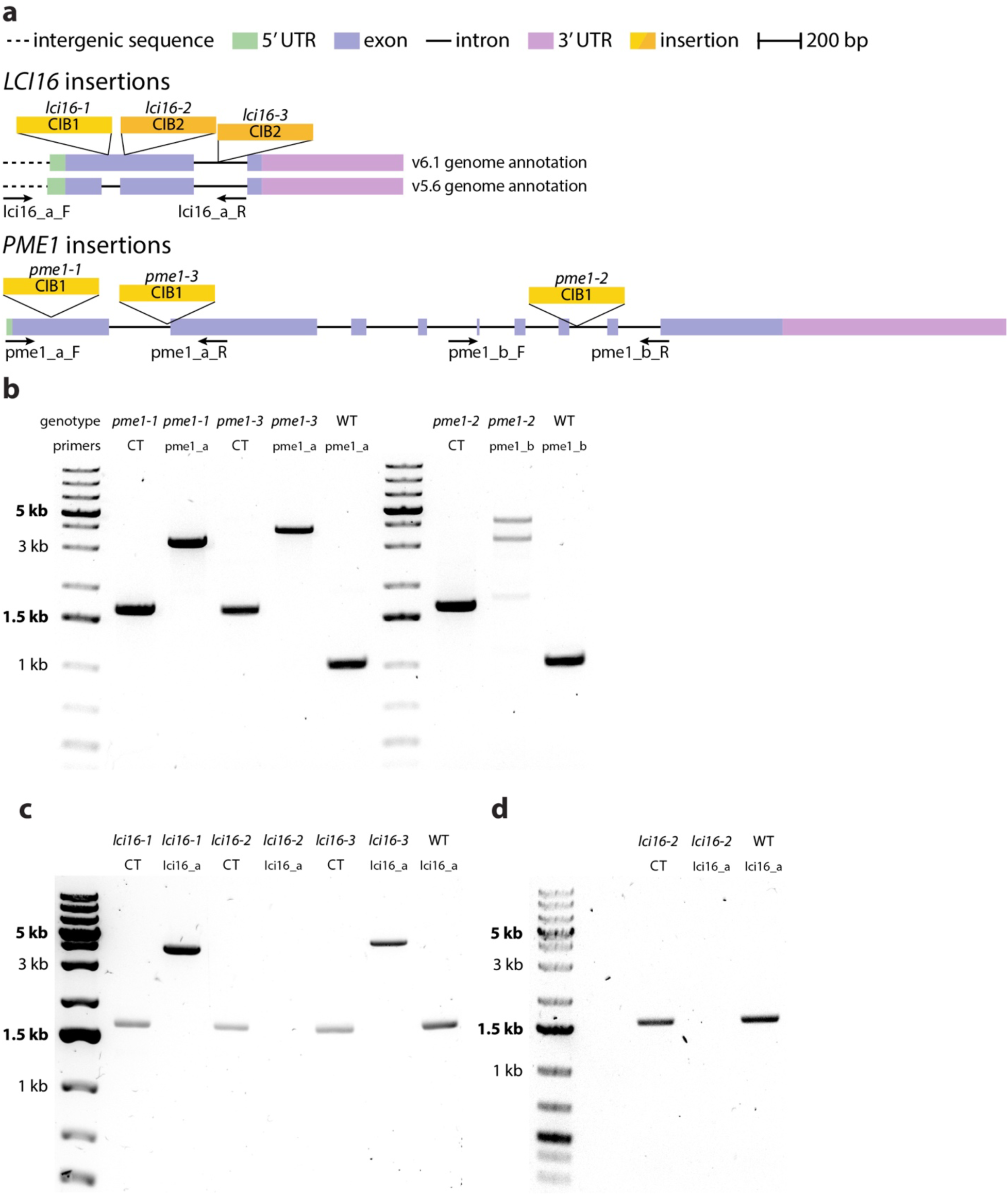
PCR verification of *lci16* and *pme1* insertional mutants. PCR verification of the insertions *lci16* and *pme1* mutants. (**a**) Maps of the LCI16 and PME1 genomic loci showing the locations of the CIB1 and CIB2 insertion cassettes and the primers used for PCR verification. The *lci16-1* insertion falls in a region that is annotated as an intron in the v5.6 *C. reinhardtii* genome assembly and an exon in the v6.1 assembly—both gene models are shown for clarity, but the insertional cassette should disrupt gene transcription regardless of whether it is inserted in an exon or intron (Li *et al*., 2019). Cassettes and primers are not drawn to scale, but distances between them are to scale. Scale bar 200 bp. (**b-d**) Agarose gels showing amplification across the insertion junction in genomic DNA purified from the *pme1* (**b**) and *lci16* (**c–d**) mutants. Expected wild-type band lengths for each primer set can be found in Supporting Information Table **S1**. For all strains except *lci16-2*, band lengths in excess of the expected wild-type amplicon lengths were observed (**b-c**), indicating the presence of the insertional cassette. The *lci16-2* gDNA failed to amplify using the insertion-spanning primers in panel (**c**), though the successful amplification using the control (CT) primers confirmed the presence of template gDNA. Panel **(d)** shows that the failure of the lci16_a primer pair to amplify *lci16-2* gDNA in independent PCR reactions is reproducible. Such a reproducible failure to amplify despite a working control is consistent with a very large insertion, and was taken as an indication of a correct insertion, as previously done (Kafri *et al*., 2023).

**Supporting Information Figure S5:**
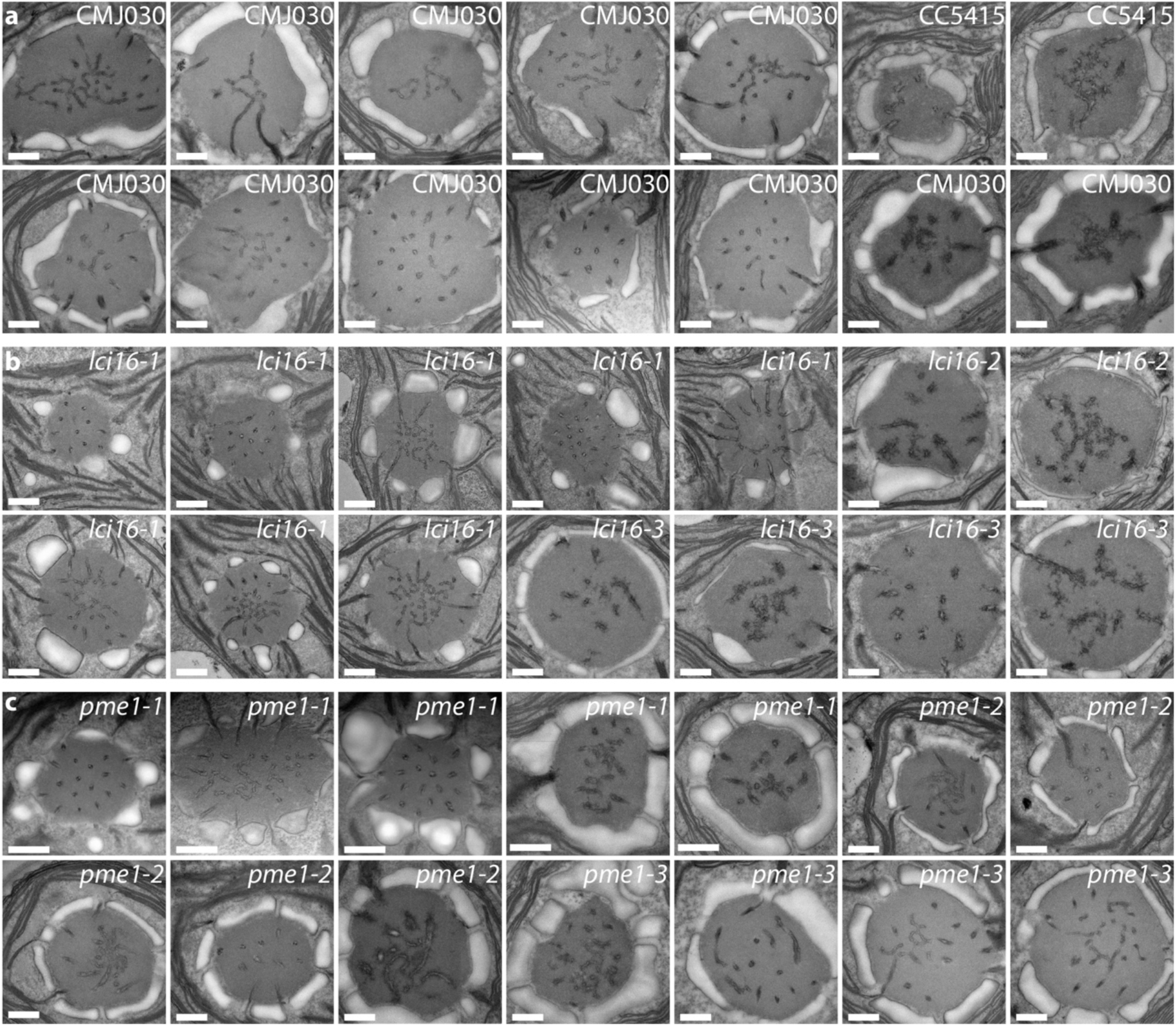
Additional TEM pyrenoid images of insertional mutants of *lci16* and *pme1*. Additional micrographs of wild-type (**a**), *lci16* (**b**), and *pme1* (**c**) pyrenoids, showing that lci16 and pme1 mutant pyrenoids contain central reticulated regions and minitubule-containing peripheral tubules. Scale bars 500 nm.

**Supporting Information Figure S6:**
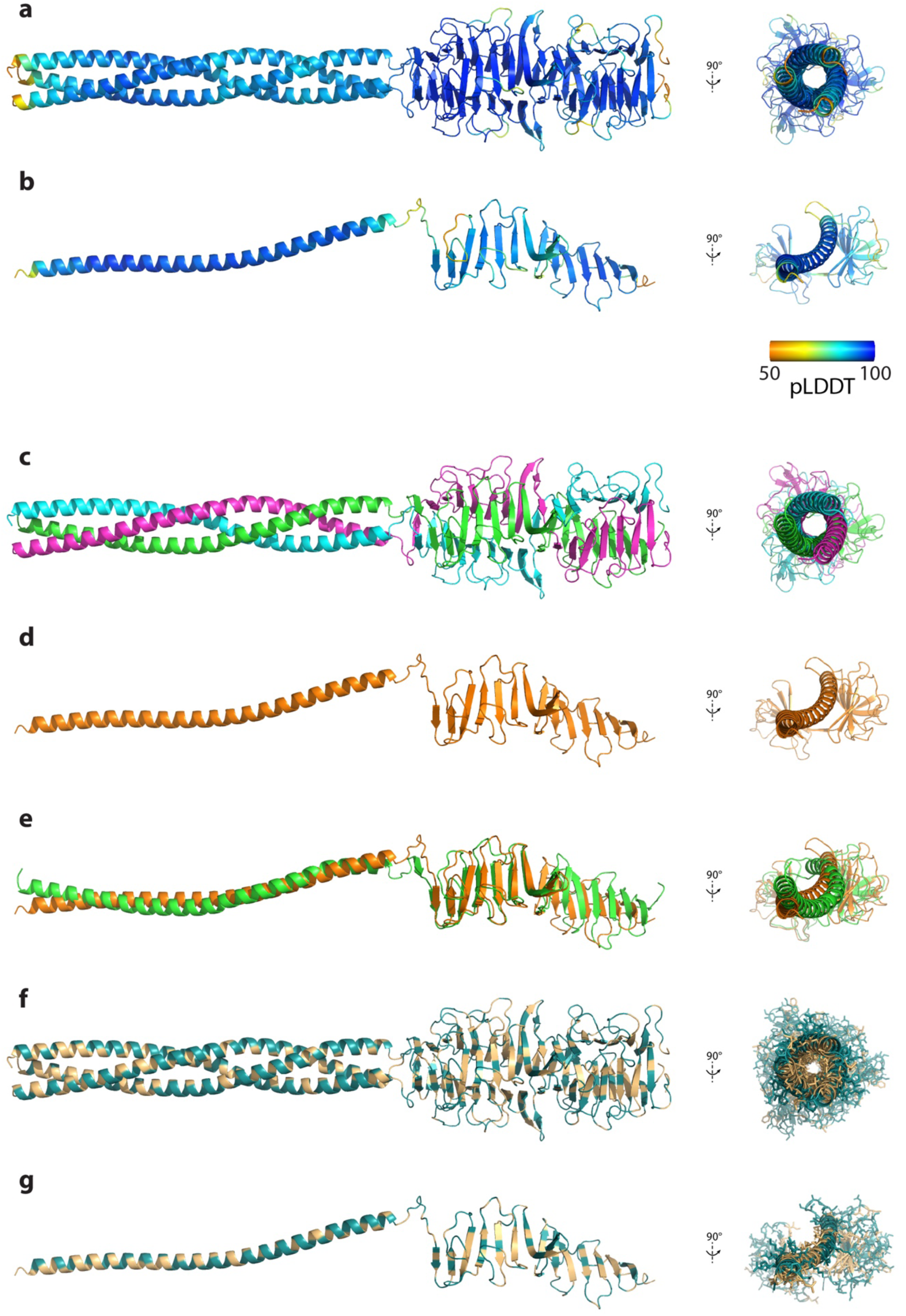
AlphaFold 3 predicts an amphipathic helix and beta sheet in PME1. AlphaFold 3 predictions of a trimer **(a,c,f)** and monomer **(b,d,g)** of residues 380–639 of PME1, the 260 residues at its C-terminus. **a–b**. The trimer **(a)** and monomer **(b)** colored according to the per-atom confidence metric pLDDT. Confidence scores for the entire structure are higher for the trimer (pTM = 0.83, iPTM = 0.82) than the monomer (pTM = 0.49). **c–e.** Cartoon representation of the extended helix and beta sheet of the predicted trimer **(c)**, in which the helices form a coiled coil and the beta sheets form a beta-barrel-like structure, and the monomer **(d)**, which takes on a similar conformation even in the absence of oligomerization, as shown by an overlay of the monomer with one subunit of the trimer **(e)**. **f–g.** The predicted trimer **(f)** and monomer **(g)** colored by Kyte-Doolittle hydrophobicity (Kyte & Doolittle, 1982), with hydrophilic residues colored deep teal and hydrophobic residues colored beige. In the 90-degree rotations showing the structure along the axis of the helix, side chain visualization has been added to show that the interior of the coiled coil contains almost exclusively hydrophobic residues, while the exterior is almost entirely hydrophilic.

**Supporting Information Table S1:**
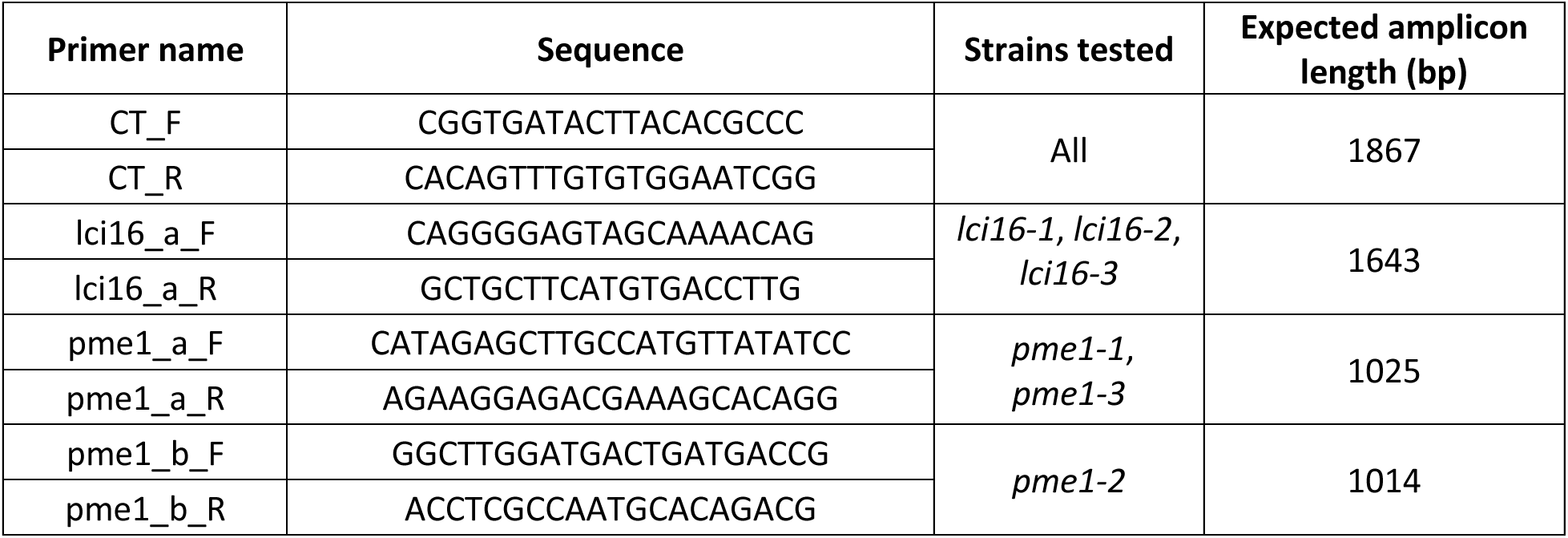
Primers used for PCR verification of *lci16* and *pme1* insertional mutants.

